# SCTC: inference of developmental potential from single-cell transcriptional complexity

**DOI:** 10.1101/2022.10.14.512265

**Authors:** Hai Lin, Huan Hu, Zhen Feng, Fei Xu, Jie Lyu, Jianwei Shuai

## Abstract

Inference of single-cell developmental potential from scRNA-Seq data enables us to reconstruct the pseudo-temporal path of cell development, which is an important and challenging task for single-cell analysis. Single-cell transcriptional diversity (SCTD), measured by the number of expressed genes per cell, has been found to be negatively correlated with the development time, and thus can be considered as a hallmark of developmental potential. However, in some cases, the gene expression level of the cells in the early stages of development may be lower than that of the later stages, which may lead to incorrect estimation of differentiation states by gene diversity-based inference. Here we refer to the economic complexity theory and propose single-cell transcriptional complexity (SCTC) metrics as a measure of single-cell developmental potential, given the intrinsic similarities between biological and economic complex systems. We take into account not only the number of genes expressed by cells, but also the more sophisticated structure information of gene expression by treating the scRNA-seq count matrix as a bipartite network. We show that complexity metrics characterize the developmental potential more accurately than the diversity metrics. Especially, in the early stages of development, cells typically have lower gene expression level than that in the later stages, while their complexity in the early stages is significantly higher than that in the later stages. Based on the measurement of SCTC, we provide an unsupervised method for accurate, robust, and transferable inference of single-cell pseudotime. Our findings suggest that the complexity emerging from the interaction between cells and genes determines the developmental potential, which may bring new insights into the understanding of biological development from the perspective of the complexity theory.

## Introduction

Single-cell RNA sequencing (scRNA-seq) technology^1,2,3^ enables a high throughput profile of gene expression for individual cells and provides new opportunities to understand the developmental process at single-cell resolution. However, due to cell’s destruction during scRNA-seq protocols, it is only possible to gain a snapshot of the cells at the collection time point, and the cells collected at this time point represent a relatively wide range of differentiation stages or cell states^4^. Therefore, how to infer single-cell developmental potential from scRNA-Seq data and then reconstruct the pseudotime of cell development is an important and challenging task in scRNA-seq studies^5,6,7,8,9^. An important research finding^10^ is that the single-cell transcriptional diversity (SCTD), measured by the number of expressed genes per cell, is a hallmark of developmental potential. That is, single-cell gene counts generally decrease with successive stages of differentiation, which provides a theoretical basis for developing a computational framework (CytoTRACE^10^) to predict differentiation states from scRNA-seq data. CytoTRACE showed good performance in most of datasets, however, under certain circumstances, such as in the earliest stage of development, cells may express fewer genes than later stages^11^. In these cases, we find that the results obtained from the method based on gene diversity cannot reflect the real developmental potential and may lead to an incorrect estimation of single-cell pseudotime. By definition, gene diversity is related only to the number of genes expressed, ignoring more subtle structural properties of gene expression, such as the ubiquity of a gene, i.e., the number of cells that express the gene. Therefore, a metrics that goes beyond gene diversity and can reflect more sophisticate structure of gene expression may be a better choice for characterizing developmental potential.

To this end, we find the complexity metrics that is used in economical area can be a candidate solution to this problem^12,13,14^. In economical context, the complexity metrics is proposed to overcome the limitation of the measure of a country’s economic development level and development potential based on the diversity of a country’s export products by adding more sophisticate productive structure information to the diversity metrics.

By analogy to economic complexity, we define single-cell transcriptional complexity (SCTC) in a similar way. In our scenario, a cell is equivalent to a country, and the genes expressed by the cell are equivalent to the products exported by the country. We regard the gene diversity, i.e., the number of genes expressed by a cell, as the 0th-order complexity of the cell, and the gene ubiquity, i.e., the number of cells that express the gene, as the 0th-order complexity of the gene. Then, by interpreting scRNA-seq data as a bipartite network in which cells are connected to the genes they express, we can define the high-order complexities of cells and genes by correcting the low-order complexities with more sophisticate network structure information. We show that the high-order complexity can better reflect the developmental potential of cells than the 0th-order complexity, especially in the initial stage of development, when cells typically have lower gene expression than subsequent stages, while their complexity is significantly higher than that in the subsequent stages, suggesting that the SCTC, which emerges from the interaction between cells and genes, is more predictive than the SCTD for cell developmental potential and is more accurate for single-cell pseudotime inference.

## Materials and Methods

### Data preparation

We calculated SCTC on four scRNA-seq datasets. The first dataset is human neuron differentiation (HND)^11^ data collected at 0, 1, 5, 7, 10 and 30 days. We removed cells with mitochondrial genes accounting for more than 15% and deleted mitochondrial genes, resulting in a dataset containing 604 cells and 13,771 genes. The second dataset is single-cell mRNA sequencing of zebrafish embryonic cells (ZEB)^15^ collected at 4, 6, 8, 10, 16, 18 and 24 hours post-fertilization (hpf), and contains 63,530 cells and 30,667 genes. The third and the fourth datasets are scRNA-seq data of human spermatogenesis (HSG) and macaque spermatogenesis (MSG)^16^, respectively. We extracted data from these two datasets for four stages of spermatogenesis, including spermatogonia, spermatocyte, round spermatid and elongating spermatid, and obtained a human spermatogenesis dataset of 10,115 cells and 45,159 genes, and a macaque spermatogenesis dataset of 19,467 cells and 22,863 genes. The selection criteria of these four datasets is based on the fact that tissue-specific genes are more abundant in brain, testis and also embryonic tissues^17^. In the following, we refer to these four datasets as HND, ZEB, HSG and MSG, respectively.

These four datasets were preprocessed based on three steps in Python package Scanpy^18,19^. Firstly we filtered out cells that do not express genes, and removed genes that are not expressed by any cells, then normalized gene expression values by Scanpy function “pp.normalize_total”, and finally performed log2 transformation on the data using Scanpy function “pp.log1p”.

### Methodology

Based on the count matrix *M*_*cg*_ of scRNA-seq data, where cell *c* express gene *g*, we define the *N*th-order complexity of cell *c* and gene *g* as^12,13^:

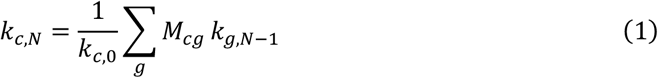

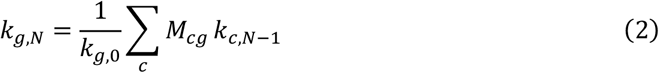

where

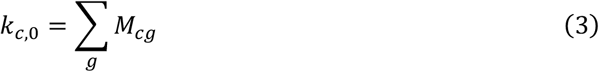

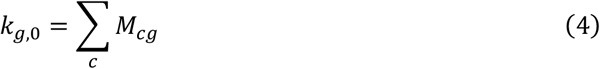

Here *k*_*c*,0_ represents the 0th-order complexity of cell *c*, which measures the diversity of the cell, and *k*_*g*,0_ is the 0th-order complexity of gene *g*, which represents the ubiquity of the gene. The high-order complexity corrects the information that low-order complexity carries by recursion. Equations (1) and (2) allow us to calculate each order complexity of cells and genes and to reconstruct pseudo-temporal path of cell development according to the complexities.

To explain concisely the theory and applications of SCTC, Fig. 1 showed a toy model in which only four cells express four genes. The scRNA-seq count matrix can be viewed as the adjacency matrix of a bipartite network, in which the gene expressions are the weights of the edges to link cells and genes (Fig. 1A). We can calculate each order of complexity recursively based on the bipartite network. Fig. 1B showed the 0th-order and 1st-order complexities of cells and genes in our toy model. Fig. 1C illustrated that the 0th-order complexity of a specific cell or a specific gene can be calculated by summing over the weights of links connected to it. Fig. 1D indicated that the higher-order complexity can be obtained by averaging over the previous order of complexity weighted by the edge weights of the bipartite network. Finally, we can sort the cells and genes according to their complexities with order *N* and infer the cell pseudotime, as shown in Fig. 1E&F.

**Fig. 1.**
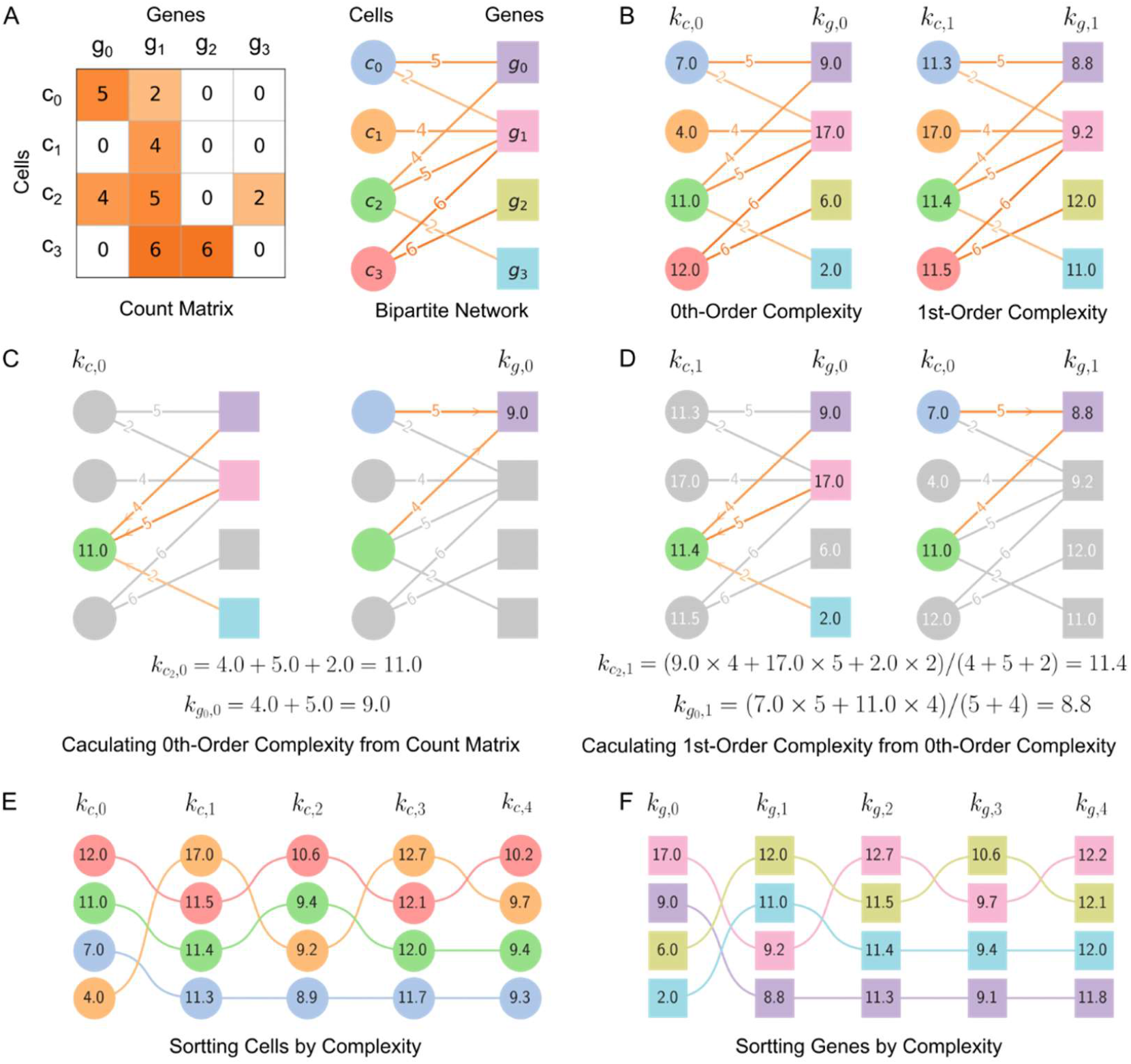
Toy model of single-cell transcriptional complexity. **A**, ScRNA-seq count matrix is viewed as the adjacency matrix of bipartite network. **B**, 0th-order and 1st-order complexities of cells and genes. **C**, Calculating 0th-order complexities of cell *c*_*2*_ and gene *g*_*0*_ by summing over the weights of edges connected to them. **D**, 1st-order complexities of cell *c*_2_ and gene g_0_ can be obtained by averaging the 0th-order complexities weighted by the weight of the edges connected to them. **E**, Sorting cells according to the cell complexity with order *N*. **F**, Sorting genes according to the gene complexity with order *N*.

As mentioned above, we can compute the complexity with each order *N* by the recursion. However, we found that, when *N* → ∞, the complexity of cells (or genes) will converge to a fixed value, which makes it difficult to choose complexity with an appropriate order *N* to characterize the developmental potential of cells. Therefore, it is necessary to find the analytic solution of the equations (1-4)^13,14^.

Substitute (2) to (1) to obtain

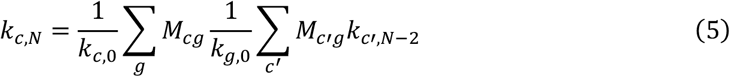

which can be rewritten as:

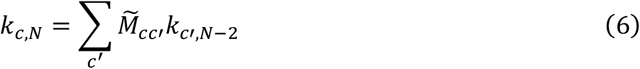

where

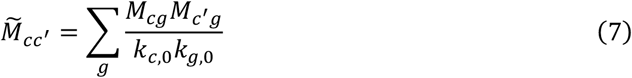

We noted that *k*_*c*,N_ = *k*_*c*,N−2_ = 1 is a solution of Eq. 6, which corresponds to the first eigenvector of 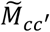. Since all components of the first eigenvector are 1 and are not informative, we take the second eigenvector 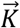 as the leading metric of cell complexity. In addition, we noted that if 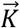 satisfies Eq.6, then 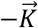 also satisfies Eq.6. Therefore, we calculated the Spearman correlation coefficient (SCC) between 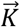 and cell diversity vector 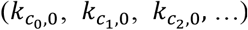, and chose the eigenvector with positive SCC. Finally, we defined the cell complexity index (CCI) as the normalized 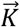:

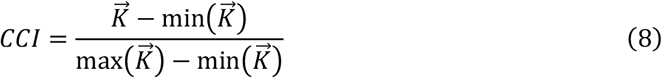

where max 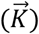 and min 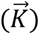 are the maximum and minimum components of 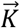, respectively.

Gene complexity index (GCI) can be defined in the same way, just by exchanging the index of cell *c* and gene *g* in the CCI definition Eq. (8). However, in order to avoid the direction problem, we defined GCI by CCI according to Eq. (2):

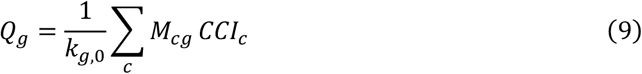

Then we defined the normalized 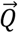 as GCI:

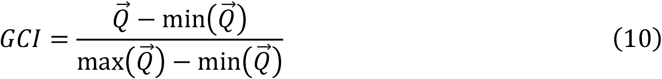

Where max 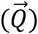 and min 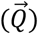 are the maximum and minimum components of 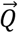, respectively.

## Results

### Comparison of pseudotime inferred by SCTD and SCTC

For four scRNA-seq data described above, we calculated gene diversity of cells, CytoTRACE^10^ pseudotime based on the SCTD, and CCI pseudotime based on the SCTC. CytoTRACE pseudotime was calculated by function “CytotraceKernel” of Python package CellRank^20^. We also used CytoTRACE R package v0.3.3^10^ to calculate the pseudotime. Although the two results obtained by CytoTRACE R package and CellRank are somehow different, it does not affect the following conclusions of this article (Supplementary Fig. S1). Since cell complexity is negatively correlated with the development time, we simply take 1-CCIc as the pseudotime of cell *c* according to Eq. (8). We compared the CytoTRACE pseudotime and the CCI pseudotime according to the time point label of cells, as shown in Fig. 2. We found that for HNC, ZEB and MSG data, cells in the early stage express fewer genes than those in the subsequent stages (top right of Fig. 2A, Fig. 2B and Fig. 2D), leading to an inaccurate inference of pseudotime by CytoTRACE (bottom left of Fig. 2A, Fig. 2B and Fig. 2D), which is consistent with the assumption that gene expression diversity is positively correlated with cell development potential^10^. In contrast, CCI pseudotime inferenced by cell complexity is in good agreement with the temporal ordering of the cells along the developmental path (bottom right of Fig. 2A, Fig. 2B and Fig. 2D), suggesting that SCTC is a better indicator of developmental potential than transcriptional diversity.

**Fig. 2.**
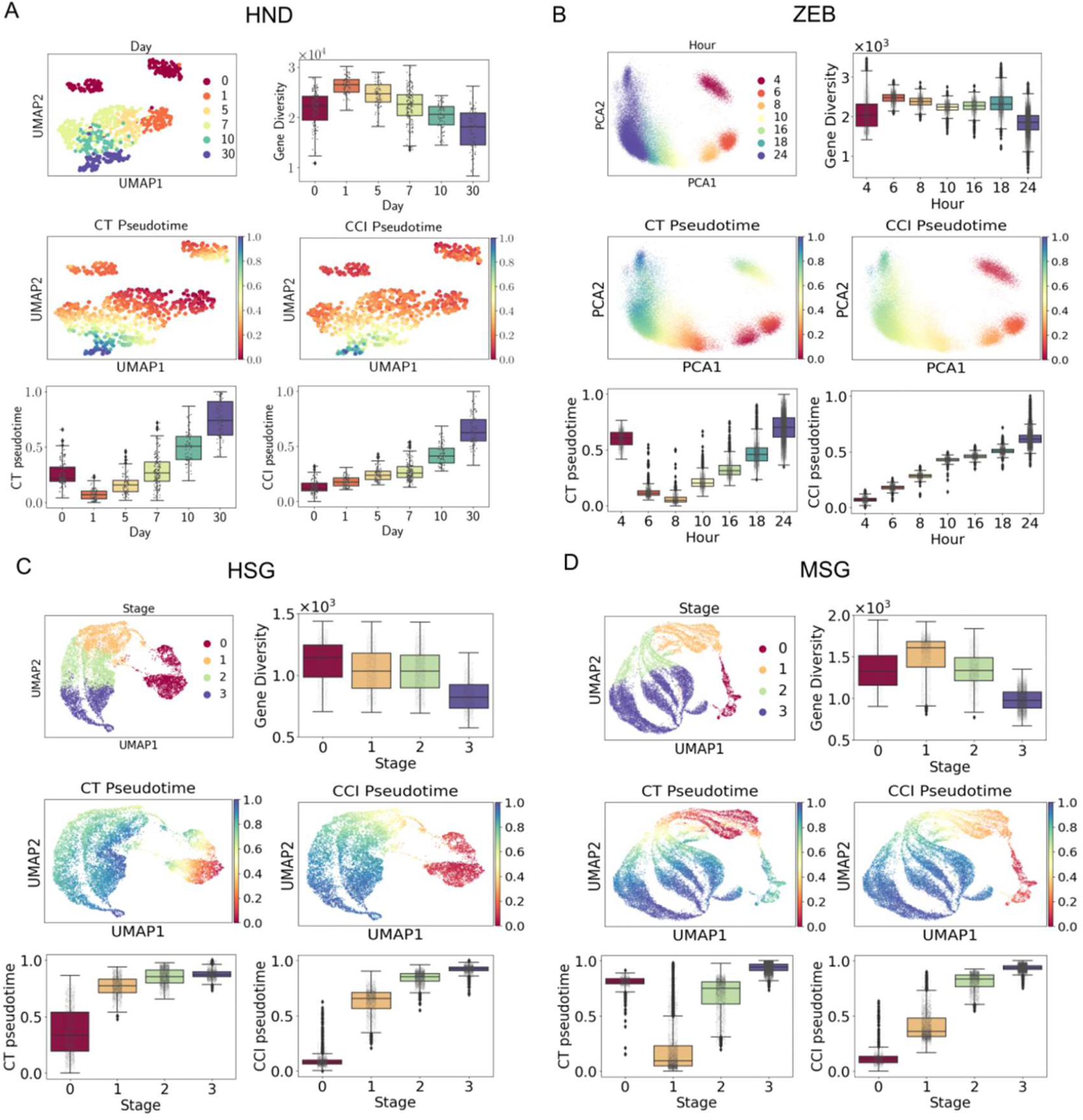
Comparison of pseudotime inferred by CytoTRACE (CT) and CCI. **A**, Human neuron differentiation (HND) data. **B**, Zebrafish embryonic cells (ZEB) data. **C**, Human spermatogenesis (HSG) data, where stage 0 stands for stage of spermatogonia, 1 for spermatocyte, 2 for round spermatid, and 3 for elongating spermatid (the same below). **D**, Macaque spermatogenesis (MSG) data. For each dataset we show UMAP (or PCA) plot of time point label (top left), box plot of gene diversity of cells at each time point (top right), UMP (or PCA) plots of CytoTRACE and CCI pseudotime (middle), and box plots of CytoTRACE and CCI pseudotime of cells at each time point (bottom), respectively.

Particularly, for the HSG data, the number of genes expressed by cells decreases monotonously with spermatogenesis process (Fig. 2C, top-right). Therefore, the pseudotime inferred by both CytoTRACE and CCI matches well with the temporal order of the development. For this dataset, we observed that the gene diversity in the first stage is only slightly higher than that in the second stage, whereas the cell complexity is much higher than that in the second stage. In addition, we found that in the first stage the heterogeneity of cell complexity is much lower than that of diversity (Fig. 2C, bottom), which implies that cell complexity metrics is more predictive of the developmental potential of cell in the early stage. Comparing Fig. 2C and Fig. 2D, we also found that the distributions of the gene diversities of HCG and MCG cells are quite different, while the distributions of cell complexity of the two species are similar, which suggested that the same tissue-of-origin genetic characteristics in different species can be efficiently identified by SCTC rather than SCTD. Taken together, the results suggest that SCTC is better than SCTD in identifying pseudotime of developmental stages in different cellular contexts.

### Explore the effect of different complexity orders in cell complexity

We recursively calculated cell complexity as a function of complexity order *N* according to equations (1-4) (Fig. 3). Since the odd-order complexities of cells are negatively correlated with the even-order complexities (Supplementary Fig. S2)^12^, we only consider the even order of cell complexity and the odd order of gene complexity in this article. Fig. 3A presented the relationship between the average cell complexity and complexity order *N* at different time points in the four scRNA-seq datasets. We found that for the development processes of the four organisms, the lower-order complexities (*N* < 8 for HND, *N* < 4 for ZEB and MSG, *N* < 2 for HSG) hardly reflect the actual developmental stages and thus fail to properly characterize the developmental potential of cells. As *N* increases, the ordering of average cell complexity tends to align well with the ordering of actual time points, suggesting that higher-order complexities are better indicator of the developmental potential of cells. However, when the order *N* exceeds a certain threshold (*N* = 16 for HND, *N* = 28 for ZEB, *N* = 52 for MSG, *N* = 60 for HSG), recursion will cause the cell complexity to collapse to the same value (Supplementary Fig. S3). To compare the numerical and analytical results, we chose an appropriate order *N*, for example, the 14th-order cell complexity, to infer the pseudotime for the four datasets and found that the CCI pseudotime calculated analytically are consistent with the results from numerical calculation (Supplementary Fig. S4), indicating that the second eigenvector of Eq. (7) accurately captures the information of cell complexity.

**Fig. 3.**
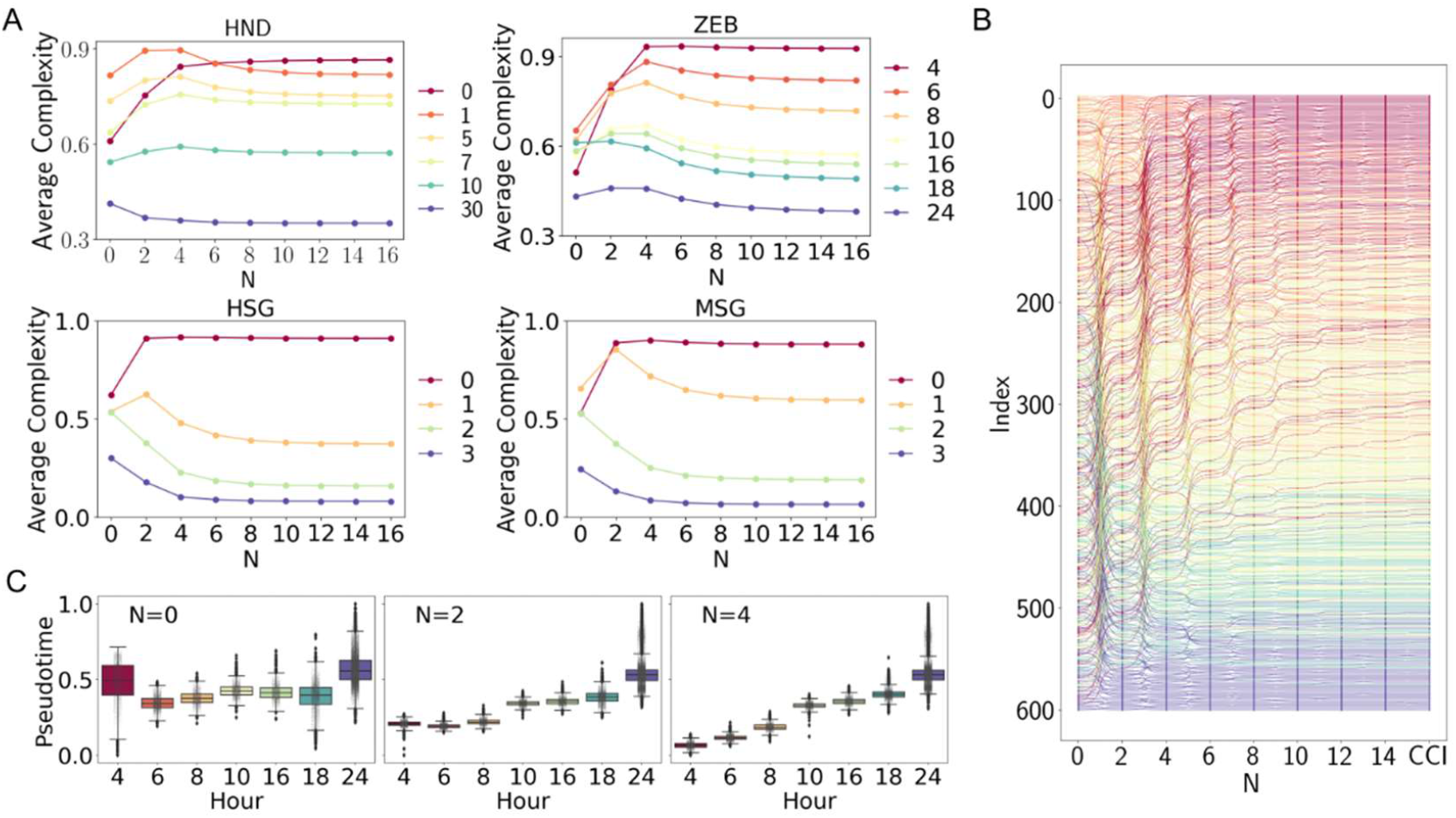
Cell complexity as a function of complexity order *N*. **A**, Average cell complexity at different time points as a function of complexity order N. **B**, 604 HND cells sorted by *N*th-order complexity and CCI. **C**, Box plots of pseudotime of ZEB cells inferred by cell complexity with different order *N*.

Figure 3B presented the results of the 604 HND cells sorted by *N*th-order complexity, with the last column showing the results sorted by CCI. The color scheme reflected different time point labels, consistent with that of the HND data in Figure 3A. It can be seen that, with the increasing complexity order N, the cells continue to rearrange and eventually converge to a stable state that is consistent with the results obtained using CCI. The rearrangement allows cells to be sorted gradually to approximate the actual time order. In Fig. 3C, we showed the pseudotime of ZEB cells inferred by cell complexity with different order *N* (*N* = 0, 2 and 4). As expected, we found that the inferred pseudotime corresponding to the 0th-order complexity is hard to explain. Instead, after four rounds of recursion, the 4th-order complexity was enough to efficiently sort cells (Fig. 3C), suggesting that the higher-order complexity captures significantly more development-relevant information than the lower-order complexity.

### The ability of gene diversity and complexity to distinguish developmental stages

As mentioned above, we referred to the number of genes expressed by a cell as the gene diversity of the cell. Similarly, we referred to the average GCI of genes expressed by a cell as the gene complexity of the cell. To evaluate the discriminative ability of the gene diversity and gene complexity in distinguishing different developmental stages, we calculated and compared the gene diversity and gene complexity of each cell in four datasets (HND, ZEB, HSG and MSG). As shown in Fig. 4A, one can see that the marginal gene diversities of cells in each stage (X-axis) have wide distributions and overlap with each other, making it difficult to distinguish different development stages. However, the marginal distributions of the gene complexity in different stages (Y-axis) are highly distinguishable, especially at the early stages where the complexity is significantly higher than those in the later stages. This phenomenon is much more pronounced in the ZEB data, where we observed that many cells are associated with much lower gene expression level in the first (4hpf) stage than in the subsequent two stages (6hpf and 8hpf), but their gene complexity is the highest during the entire 24 hours of development (Fig. 4A). Altogether, the single-cell transcriptional complexity is a more accurate indicator of developmental potential than diversity. At the final stage of development, most of the cells fall into the third quadrant formed by two dashed lines (FIG. 4A), indicating that both gene diversity and gene complexity are at a low level for the differentiated cells, implying a looser gene regulation relationship in later stages than earlier stages.

**Fig. 4.**
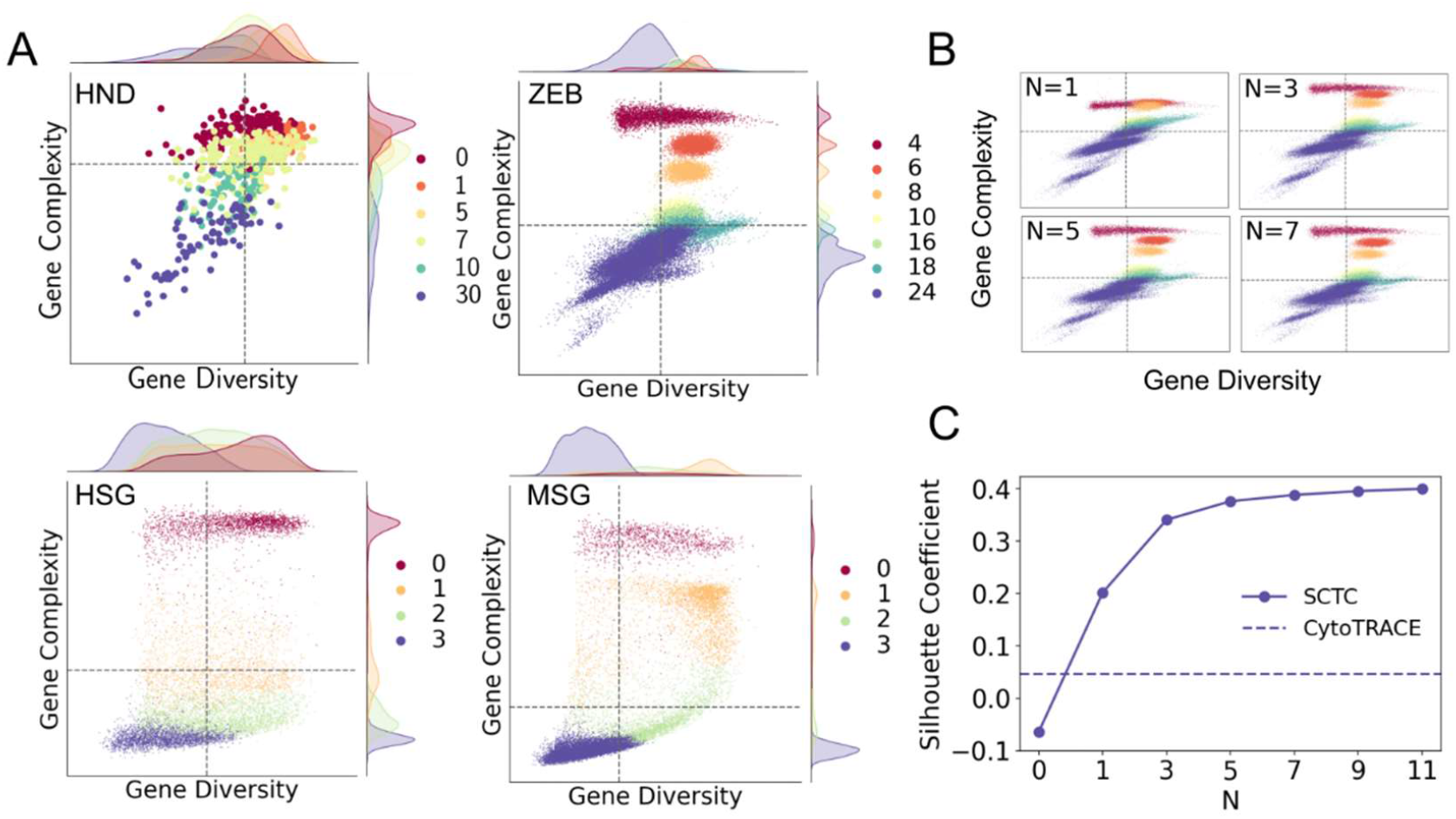
The ability of gene diversity and complexity to distinguish developmental stages. **A**, The diversity-complexity diagrams of single-cell gene expression. Dash lines indicate the mean diversity and mean complexity averaged over all cells. **B**, The diversity-complexity diagram as a function of gene complexity order *N* for the ZEB data. **C**, Silhouette coefficient of gene complexity as a function of complexity order *N* for the ZEB data, where *N* = 0 denoted the gene diversity. Dash line indicated the Silhouette coefficient of CytoTRACE pseudotime.

We also investigated the relationship between gene diversity and gene complexity with order *N* for the ZEB data (Fig. 4B). We found that gene complexity can distinguish different cell differentiation stages very well, especially at higher gene complexity order *N* (e.g., *N* = 5 or 7). To make it clear, we calculated the Silhouette coefficient^21^, a metrics measures how well the clusters are apart from each other and clearly distinguished, of the gene complexity according to the reverse order of time point label as a function of complexity order *N*. Fig. 4C showed the results of the ZEB data, where *N* = 0 denote gene diversity. For comparison, Silhouette coefficient of CytoTRACE pseudotime was also calculated. We found that gene complexity has a higher Silhouette coefficient than gene diversity, and the Silhouette coefficient increases with the increasing of complexity order N. Similar trends were also observed in the other three datasets (Supplementary Fig. S5), indicating that the higher-order gene complexity has a better ability to distinguish developmental stages than low-order gene complexity or gene diversity.

### Genes of different GCI are associated with specific developmental stages

We randomly selected 600 genes in the HND dataset to study how they are sorted in terms of complexity with different order N (Fig. 5A). Since there was no experimental measurement of gene complexity, we used GCI defined by Eq. (10) as a benchmark to investigate the process of gene rearrangement according to the complexity with different order N. As shown in Fig. 5A, the ordering of genes was adjusted continuously as the order N increases, with some genes undergoing a large span of rearrangement and eventually converging to a stable stage that was consistent with the ordering of GCI.

**Fig. 5.**
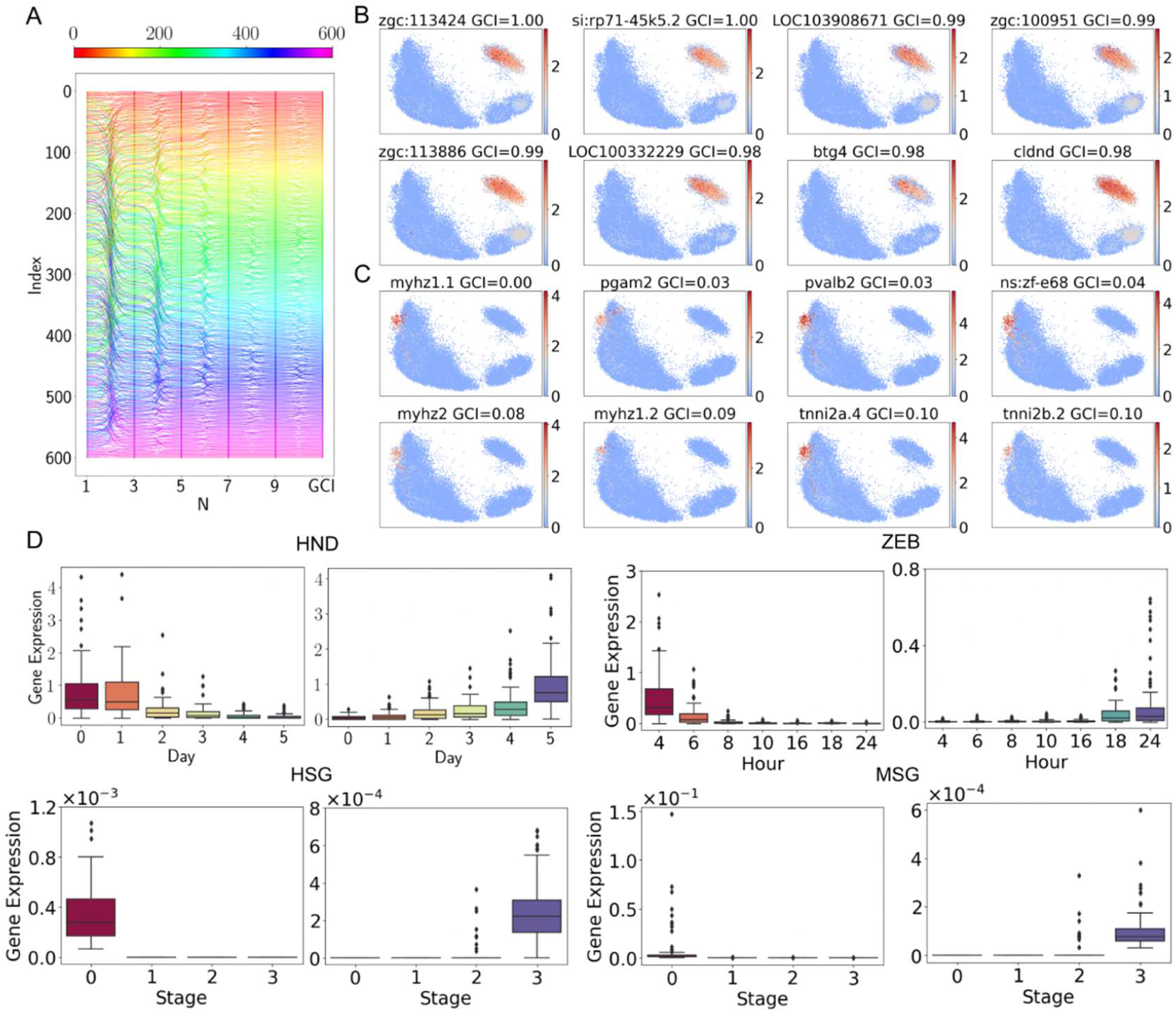
Association of genes of different GCI with developmental stages. **A**, 600 randomly selected HND genes sorted by *N*th-order complexity and GCI. **B**, Distribution of top eight genes in ZEB data. **C**, Distribution of bottom eight genes in ZEB data. **D**. top 100 higher-GCI genes (left in each panel) and bottom 100 lower-GCI genes (right in each panel) at each time points.

We then examined the developmental stage distributions of genes with top ranking and bottom ranking in complexity. Fig. 5B and Fig. 5C showed the distributions of top eight genes and bottom eight genes in ZEB data, from which we observed that the genes with the highest complexity are mainly expressed at the first time point of development, whereas the genes with the lowest complexity are mainly distributed at the final time point. Similar trends were also observed in the other three datasets (Supplementary Fig. S5). Furthermore, we analyzed the gene expression levels of top 100 higher-GCI genes and bottom 100 lower-GCI genes at each time points for the four datasets (Fig. 5D). Interestingly, we observed that the genes with high complexity are mainly expressed in the earlier stages of development, whereas the genes with low complexity are mainly expressed in the later stages, suggesting a strong correlation between gene complexity as measured by GCI and cell development potential (Fig. 5D). Taken together, we associated genes expressed in different cell developmental stages with higher or lower GCI, which may also be useful for annotating cell development related genes.

### Transferability of single-cell transcriptional complexity model

In our method, cell complexity and gene complexity are recursively defined by each other as indicated by equations (1) and (2), therefore, the cell complexity can be computed from gene complexity and vice versa. In addition, many genes are presented in more than one single-cell dataset, therefore, gene complexity calculated by one dataset can be used to calculate cell complexity of another dataset, in another word, our model is transferable in practice.

To this end, we first merged the HSG data and MSG data according to the genes shared by the two datasets, and obtained a mixed dataset with 29,591 cells and 14,405 genes. Fig. 6A showed the UMAP plot of the development stages of HSG and MSG cells, and Fig. 6B showed the distribution of the gene diversity of the cells in merged dataset. We calculated CytoTRACE pseudotime and CCI pseudotime of this merged dataset, as shown in Fig. 6C. It can be seen that the distribution of CytoTRACE pseudotime is relatively disordered, whereas the CCI pseudotime maintains the correct ordering of the two types of cells, indicating that the SCTC model is more robust to the mixed datasets than the SCTD model.

**Fig. 6.**
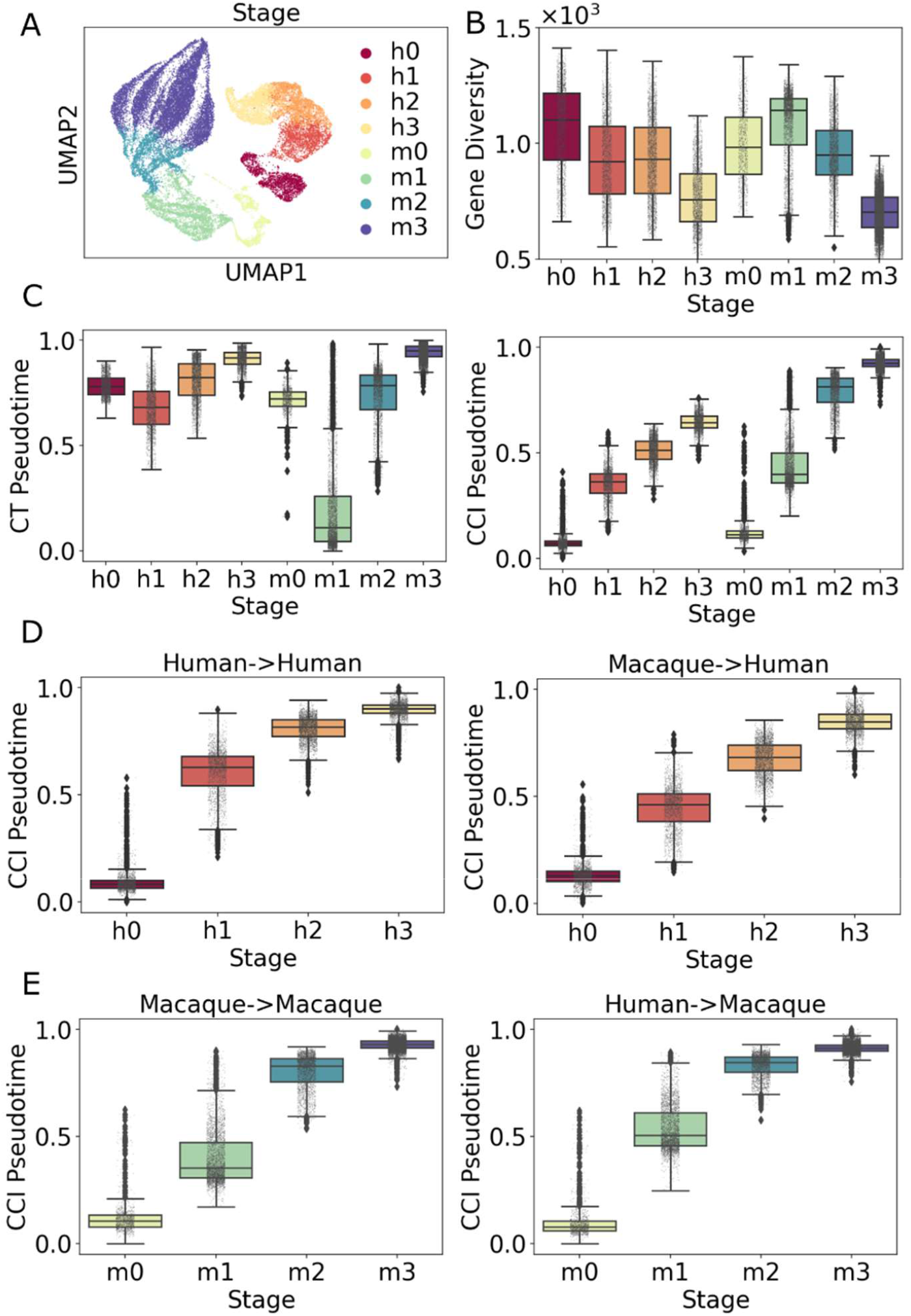
Transferability of single-cell transcriptional complexity model. **A**, UMAP plot of the development stage of the merged dataset. “h” denotes human and “m” denotes macaque (the same below). **B**, Box plot of gene diversity of cells in merged dataset. **C**, Box plots of CytoTRACE pseudotime (left) and CCI pseudotime (right) of merged dataset. **D**, Human CCI pseudotime computed by human GCI (left) and by macaque GCI (right). **E**, Macaque CCI pseudotime computed by macaque GCI (left) and by human GCI (right).

In order to examine the transferability of complexity model, we then divide the merged dataset into two new datasets according to human cells and macaque cells, namely the new HCG data and the new MCG data, which shared 14,405 genes in common. We first calculated the GCIs for the two new datasets. Then, according to Eq. (1), the human and macaque GCIs were used to calculate the human CCI pseudotime (Fig. 6D), and the macaque and human GCIs were used to calculate the macaque CCI pseudotime (Fig. 6E). Figs. 6D&E showed that the CCI pseudotime calculated with GCI of other datasets was not much different from that calculated with GCI of their own dataset, indicating that the complexity model has good transferability between different datasets. The results further demonstrated the conservation of gene complexity in spermatogenesis among different species, and provide a new insight of the application of complexity theory in the inference of cell developmental potential in different species.

## Discussion

The unprecedented resolution of single-cell RNA-sequencing data provides challenges in dissecting the underlying biological mechanisms. In this work, we introduce single-cell transcriptional complexity to infer pseudotime and the developmental potential and find it is superior to available methods in recovering developmental processes such as neuronal development and spermatogenesis. The proposed method, single-cell transcriptional complexity is inspired from the economic complexity theory that was used in the economy area, which was used to evaluate the development level and developmental potential of countries in the economic system. When evaluating the complexity theory in cellular context, we find that in the early development stage, though cells may have lower gene expression level than later stages, their complexity with appropriate order *N* is significantly higher than that in the later stages, indicating that complexity is a better indicator of the developmental potential of cells than diversity. The finding can be explained that a cooperation of pluripotency genes in a large interconnected network rather than gene expression intensity of pluripotency related genes can determine the developmental potential^22^, which can be captured by transcriptional complexity metric that we propose.

Based on transcriptional complexity, we provide a simple yet efficient method for single-cell pseudotime inference, which relies on the scRNA-seq count matrix while without a need to choose any highly variable genes^20^. The obtained pseudotime matches the actual time point labels of cells more accurately than the diversity-based method, for example, CytoTRACE (Fig. 2). We find the performance of CytoTRACE method in certain datasets is worse than our method, particularly in the earliest stage of development where cells may express fewer genes than later stages, which can be explained by the underlying diversity-based assumption of CytoTRACE does not hold true in this scenario. What is more, our method is also robust to the merged dataset of different data and is transferable between different datasets (Fig. 6), which makes it possible to further investigate the meaning of cell and gene complexity in different organisms or in biological contexts.

Actually, there are some dedicated computational tools have been developed for the developmental potential inference of single-cell sequencing data except for CytoTRACE^6,23,24,25^. The state-of-the-art methods are CCAT^24^ and FitDevo^25^. However, both methods are supervised. CCAT needs prior knowledge, and FitDevo relies on the training set, which may limit their scope of application, as can be seen in supplementary Fig. S7, where we show the pseudotime of four datasets inferred by FitDevo and SCTC, respectively. We found that for the HND data, which is one of the training datasets in FitDevo, both methods achieve a high SCC score, while for other three datasets not included in the training datasets of FitDevo, SCTC performs much better than FitDevo despite being an unsupervised method. Especially for the ZEB data, which is removed from the training set in FitDevo due to the limited number of homologous genes between zebrafish and mammals^25^, FitDevo made an incorrect estimation of developmental potential at early developmental stages as CytoTRACE does. Therefore, an unsupervised inference method like SCTC may be a better choice in this scenario.

Our method borrows the method of economic complex systems to the single cell transcriptional analysis and obtains meaningful results, indicating that there are some intrinsic similarities between biological and economic complex systems. Alternatively, a series of other theories and technologies which have been developed in the field of economic complex systems^14,26,27,28^ may also be applied to the field of single-cell study and beyond, which may provide new insights for understanding biological development from the perspective of complex systems^29,30^.

## Data and Code Availability

The source code and the data of filtered human neuron differentiation (HND) are available at https://github.com/hailinphysics/sctc. The raw data of HND^11^ can be accessed from Gene Expression Omnibus (GEO) through the accession number GSE102066. Zebrafish embryonic cells (ZEB)^15^ dataset can be accessed from GEO under accession number GSE112294. Human and macaque spermatogenesis datasets^16^ are available under the GEO accession number GSE142585.

## References

1. Tang, F. et al. mRNA-Seq whole-transcriptome analysis of a single cell. Nat. Methods 6, 377–382 (2009).

2. Parekh, S. et al. Comparative Analysis of Single-Cell RNA Sequencing Methods. Mol. Cell 65, 631–643 (2017).

3. Hwang, B., Lee, J. H. & Bang, D. Single-cell RNA sequencing technologies and bioinformatics pipelines. Exp. Mol. Med. 50, 1–14 (2018).

4. Ding, J., Sharon, N. & Bar-Joseph, Z. Temporal modelling using single-cell transcriptomics. Nat. Rev. Genet. 23, 355–368 (2022).

5. Ji, Z. & Ji, H. TSCAN: Pseudo-time reconstruction and evaluation in single-cell RNA-seq analysis. Nucleic Acids Res. 44, e117 (2016).

6. Teschendorff, A. E. & Enver, T. Single-cell entropy for accurate estimation of differentiation potency from a cell’s transcriptome. Nat. Commun. 8, 1–15 (2017).

7. Street, K., Risso, D., Fletcher, R. et al. Slingshot: cell lineage and pseudotime inference for single-cell transcriptomics. BMC Genomics 19, 477 (2018).

8. Mealy, P., Farmer, J. D. & Teytelboym, A. Interpreting economic complexity. Sci. Adv. 14, 1–9 (2019).

9. Saelens, W., Cannoodt, R., Todorov, H. & Saeys, Y. A comparison of single-cell trajectory inference methods. Nat. Biotechnol. 37, 547–554 (2019).

10. Gulati, G. S. et al. Single-cell transcriptional diversity is a hallmark of developmental potential. Science. 367, 405–411 (2020).

11. Wang, J. et al. Single-cell gene expression analysis reveals regulators of distinct cell subpopulations among developing human neurons. Genome Res. 27, 1783–1794 (2017).

12. Hidalgo, C. A. & Hausmann, R. The building blocks of economic complexity. Proc. Natl. Acad. Sci. U. S. A. 106, 10570–10575 (2009).

13. Hausmann, R. et al. The Atlas of Economic Complexity: Mapping Paths to Prosperity. (The MIT Press, 2014).

14. Hidalgo, C. A. Economic complexity theory and applications. Nat. Rev. Phys. 3, 92– 113 (2021).

15. Wagner, D. E. et al. Single-cell mapping of gene expression landscapes and lineage in the zebrafish embryo. Science. 360, 981–987 (2018).

16. Shami, A. N. et al. Single-Cell RNA Sequencing of Human, Macaque, and Mouse Testes Uncovers Conserved and Divergent Features of Mammalian Spermatogenesis. Dev. Cell 54, 529-547.e12 (2020).

17. Lin, S. et al. Comparison of the transcriptional landscapes between human and mouse tissues. Proc. Natl. Acad. Sci. U. S. A. 111, 17224–17229 (2014).

18. Wolf, F. A., Angerer, P. & Theis, F. J. SCANPY: Large-scale single-cell gene expression data analysis. Genome Biol. 19, 15 (2018).

19. Luecken, M. D. & Theis, F. J. Current best practices in single-cell RNA-seq analysis: a tutorial. Mol. Syst. Biol. 15, (2019).

20. Lange, M. et al. CellRank for directed single-cell fate mapping. Nature Methods vol. 19 (2022).

21. Rousseeuw, P. J. Silhouettes: A graphical aid to the interpretation and validation of cluster analysis. J. Comput. Appl. Math. 20, 53–65 (1987).

22. Li, M. & Izpisua Belmonte, J. C. Deconstructing the pluripotency gene regulatory network. Nat. Cell Biol. 20, 382–392 (2018).

23. Guo, M., Bao, E. L., Wagner, M., Whitsett, J. A. & Xu, Y. SLICE: Determining cell differentiation and lineage based on single cell entropy. Nucleic Acids Res. 45, 1–14 (2017).

24. Teschendorff, A. E., Maity, A. K., Hu, X., Weiyan, C. & Lechner, M. Ultra-fast scalable estimation of single-cell differentiation potency from scRNA-Seq data. Bioinformatics 37, 1528–1534 (2021).

25. Zhang, F. et al. FitDevo: accurate inference of single-cell developmental potential using sample-specific gene weight. Brief. Bioinform. 23, 1–13 (2022).

26. Hidalgo, C. A., Winger, B., Barabási, A. L. & Hausmann, R. The product space conditions the development of nations. Science. 317, 482–487 (2007).

27. Caldarelli, G. et al. A Network Analysis of Countries’ Export Flows: Firm Grounds for the Building Blocks of the Economy. PLoS One 7, 1–11 (2012).

28. Hynes, W., Trump, B. D., Kirman, A., Haldane, A. & Linkov, I. Systemic resilience in economics. Nat. Rev. Phys. 18, 381–384 (2022).

29. Macarthur, B. D. & Lemischka, I. R. Statistical mechanics of pluripotency. Cell 154, 484–489 (2013).

30. Teschendorff, A. E. & Feinberg, A. P. Statistical mechanics meets single-cell biology. Nat. Rev. Genet. 22, 459–476 (2021).

