## Supplementary Materials for "SCTC: inference of developmental potential from single-cell transcriptional complexity"

### Supplementary Materials for
### SCTC: inference of developmental potential from single-cell transcriptional complexity

Hai Lin[1,2], Huan Hu[3], Zhen Feng[4], Fei Xu[3], Jie Lyu[2,5], Jianwei Shuai[1,2,3,6,*]

[1]Wenzhou Key Laboratory of Biophysics, Wenzhou Institute, University of Chinese Academy of Sciences, Wenzhou, Zhejiang 325001, China

[2]Oujiang Laboratory (Zhejiang Lab for Regenerative Medicine, Vision and Brain Health), Wenzhou, Zhejiang 325001, China

[3]Department of Physics, and Fujian Provincial Key Laboratory for Soft Functional Materials Research, Xiamen University, Xiamen 361005, China

[4]First Affiliated Hospital of Wenzhou Medical University, Wenzhou Medical University, Wenzhou 325000, China

[5]Joint Centre of Translational Medicine, Wenzhou Institute, University of Chinese Academy of Sciences, Wenzhou, Zhejiang 325001, China

[6]National Institute for Data Science in Health and Medicine, and State Key Laboratory of Cellular Stress Biology, Innovation Center for Cell Signaling Network, Xiamen University, Xiamen 361102, China

[*]

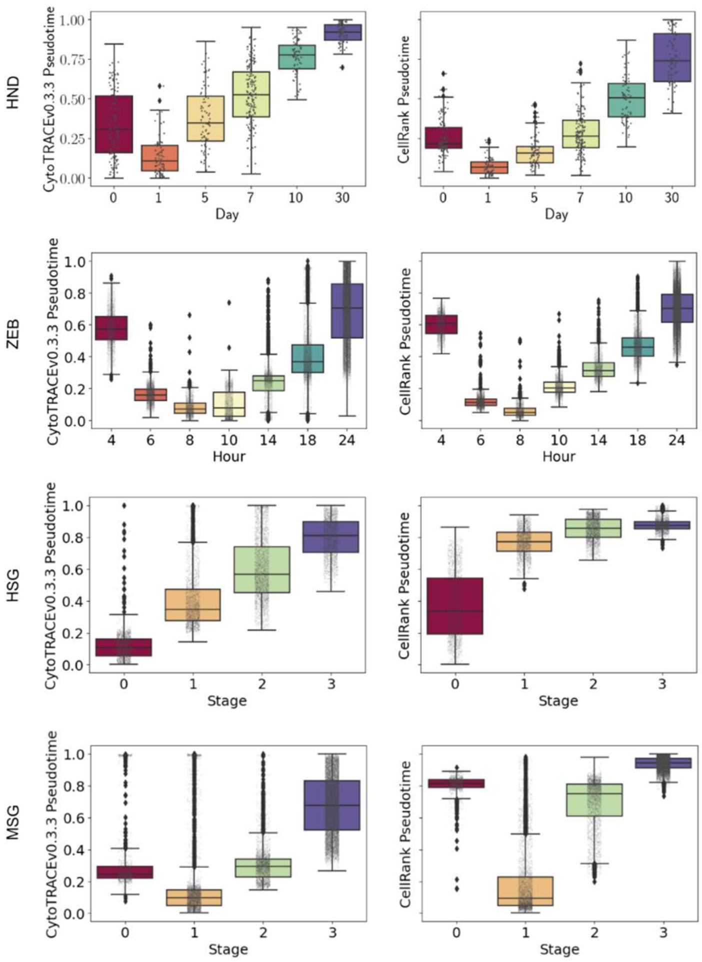


**Fig. S1,** Comparison of the pseudotime calculated by CellRank CytotraceKernel (left) and by CytoTRACE R package v0.3.3 (right).

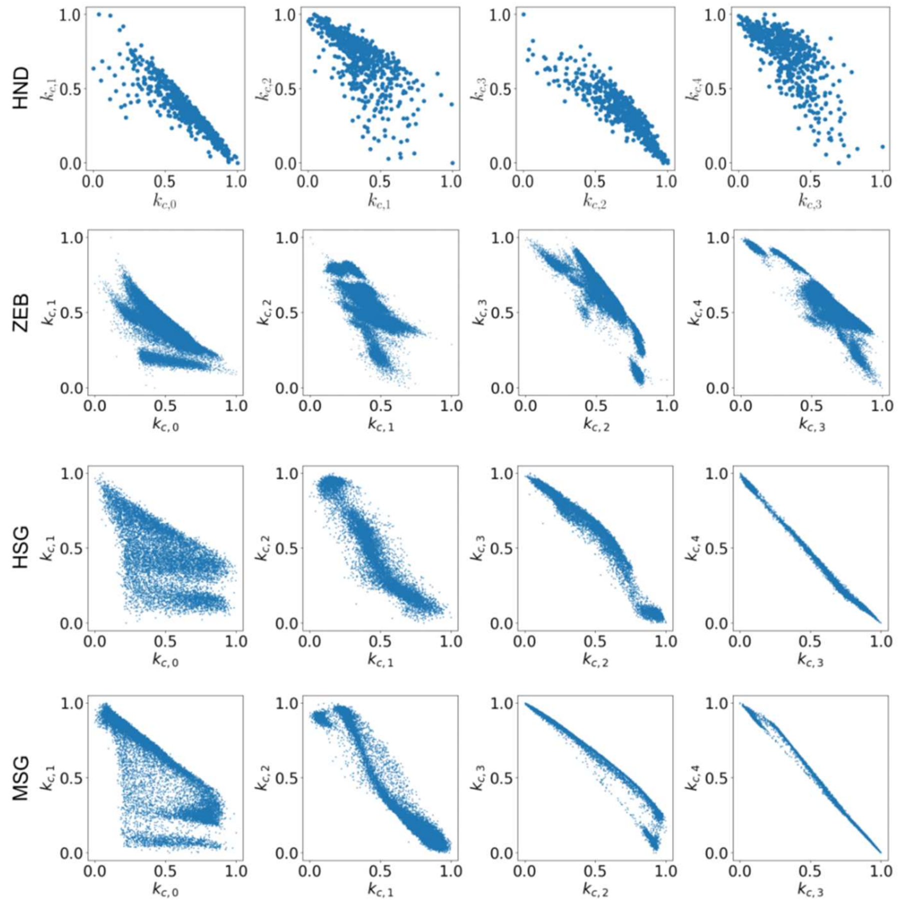


**Fig. S2,** The odd-order complexity of cells is negatively correlated with the even-order complexity.

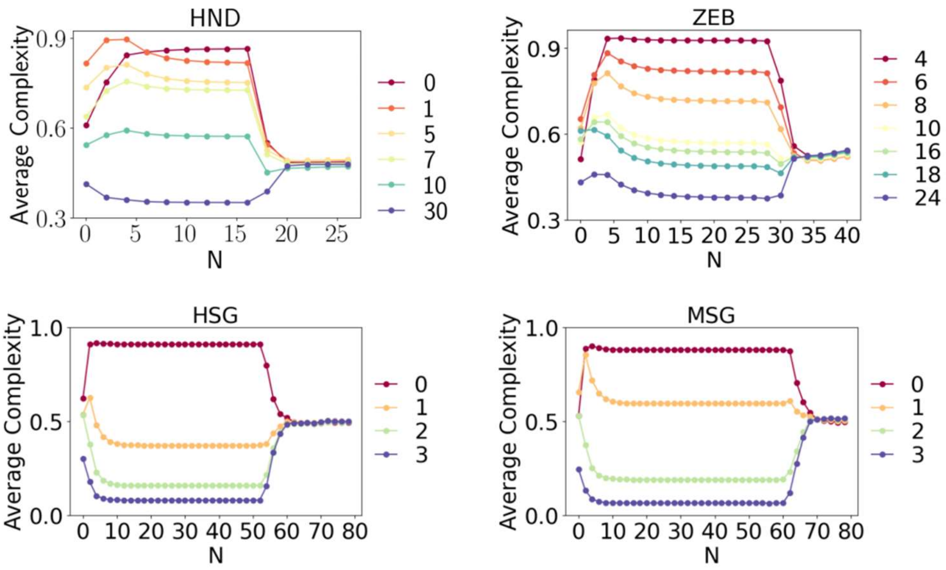


**Fig. S3,** When the complexity order N exceeds a certain threshold, recursion will cause the cell complexity to collapse to the same value.

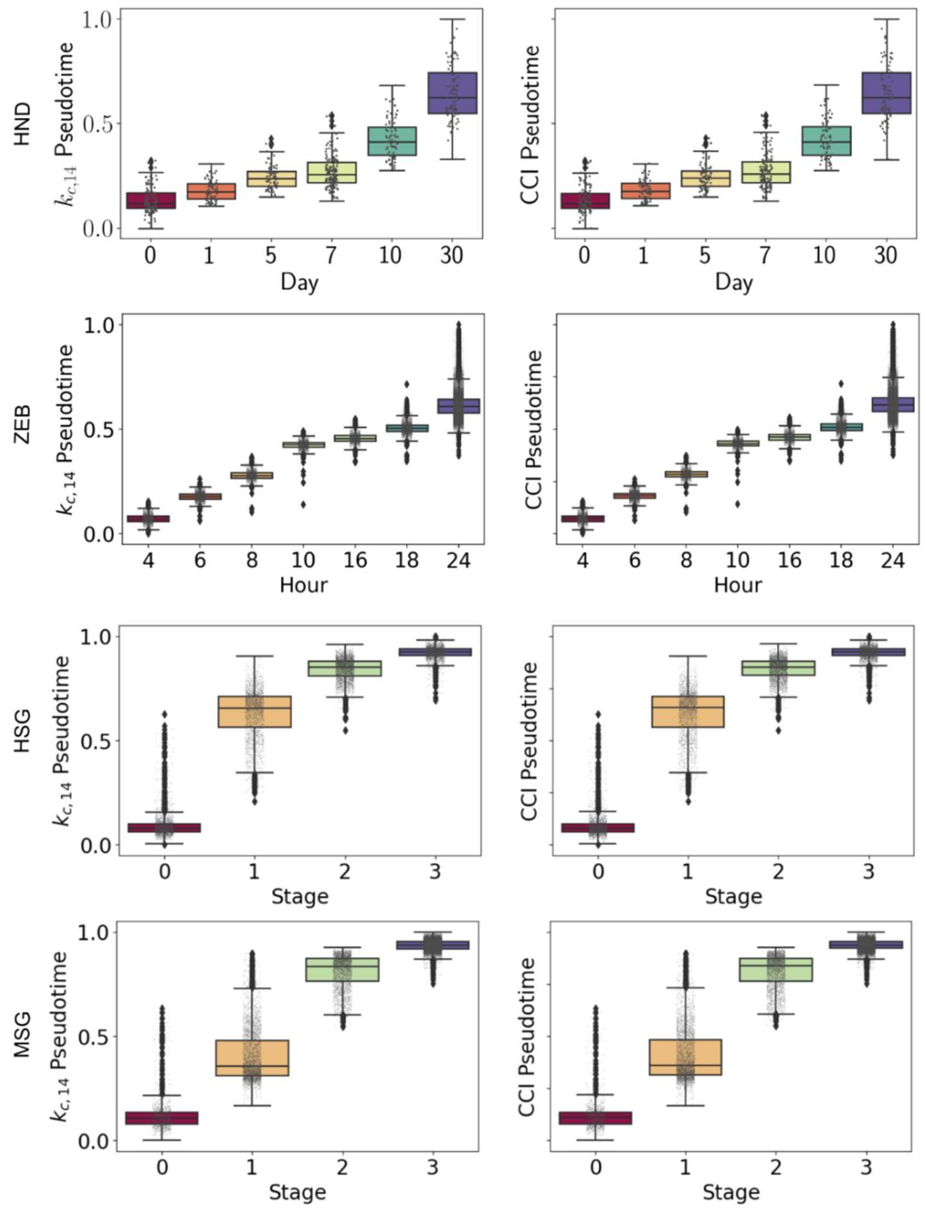


**Fig. S4,** The pseudotime inferred by $k_{c,14}$ (left) are no different from that inferred by CCI (right).

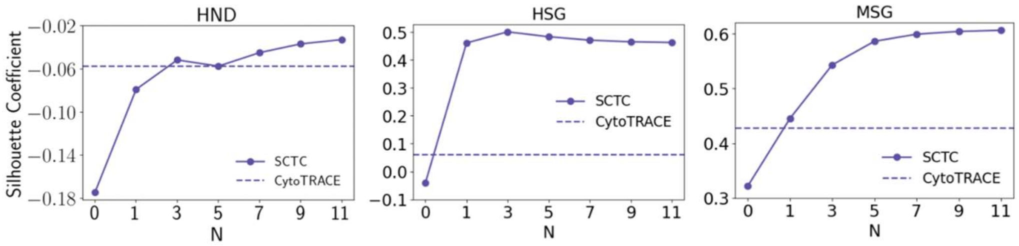


**Fig.S5**, Silhouette coefficient of gene complexity as a function of complexity order $N$ for HND data, HSG data and MSG data.

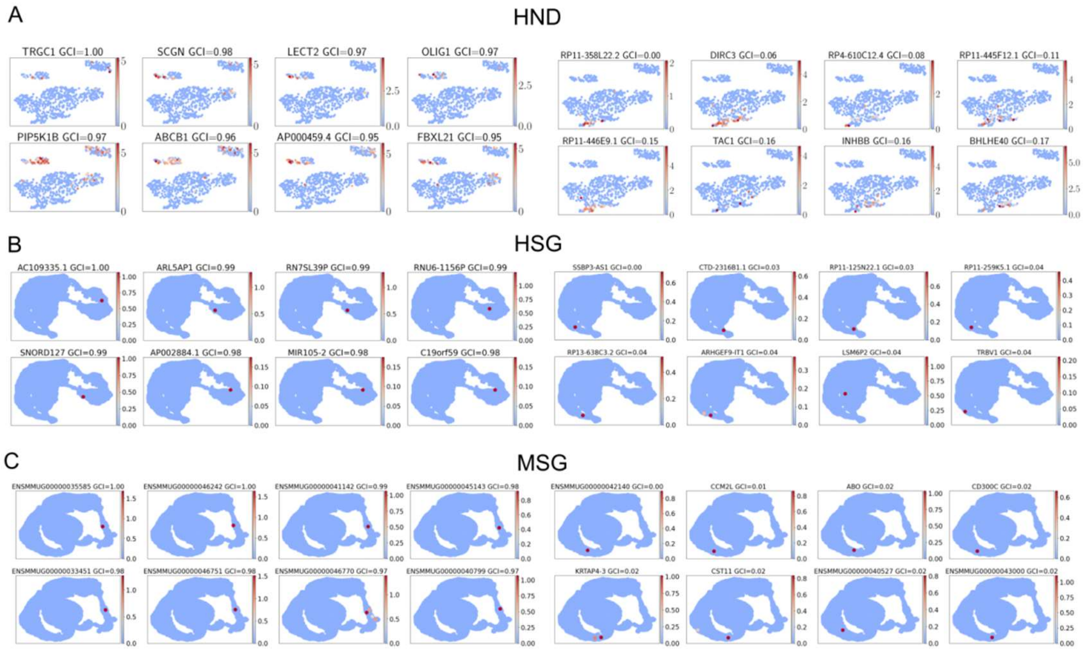


**Fig. S6,** Distribution of top eight genes (left) and bottom eight genes (right) for **A**, HND data, **B**, HSG data and **C**, MSG data.

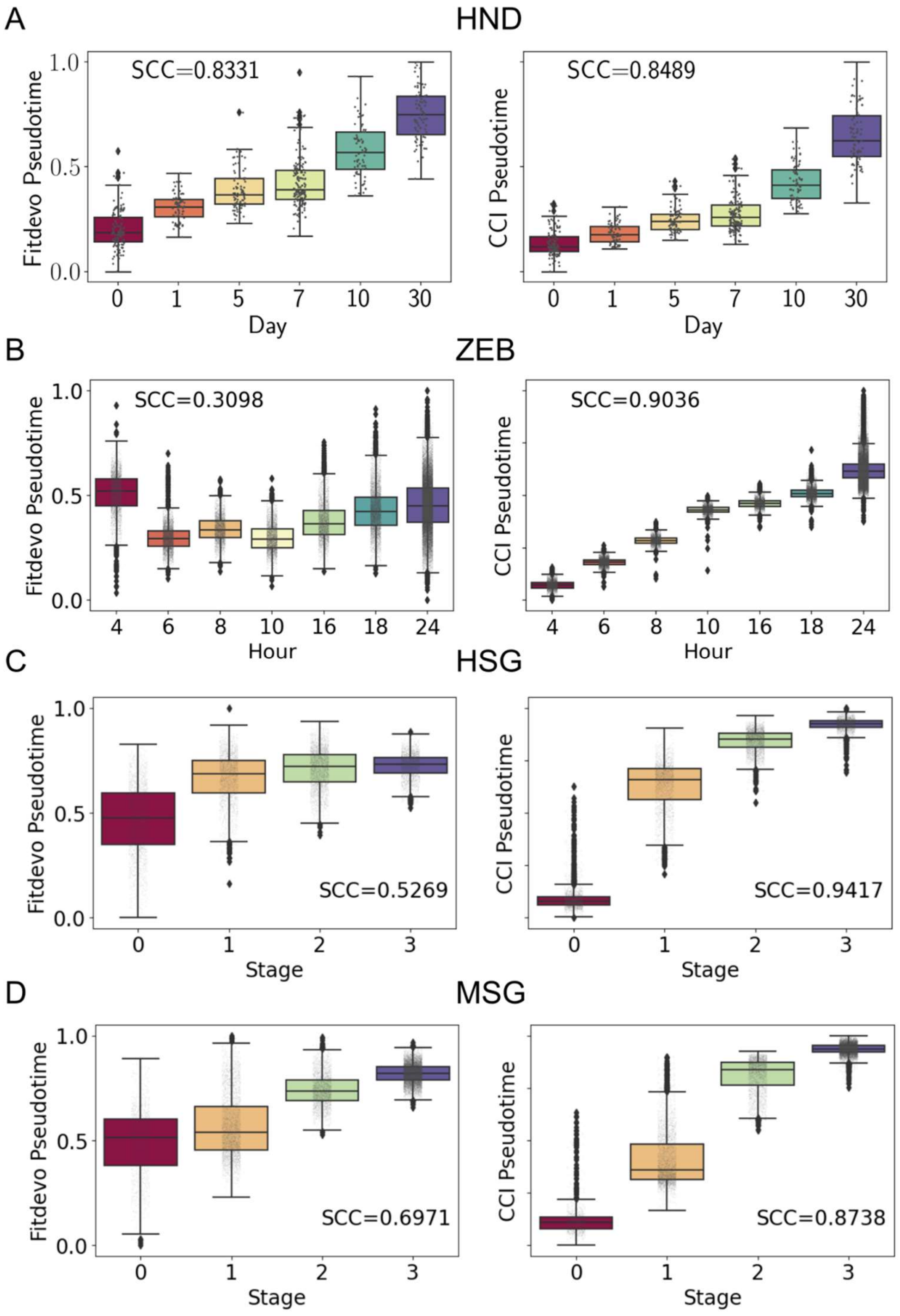


**Fig.S7**，Comparison of pseudotime inferred by FitDevo (left) and SCTC (right) for **A,** HND data, **B,** ZEB data, **C,** HSG data and **D,** MSG data.
